# Identifying small molecule binding sites for epigenetic proteins at domain-domain interfaces

**DOI:** 10.1101/283069

**Authors:** David Bowkett, Romain Talon, Cynthia Tallant, Chris Schofield, Frank von Delft, Stefan Knapp, Gordon Bruton, Paul E. Brennan

## Abstract

Epigenetics is of rapidly growing field in drug discovery. Of particular interest is the role of post-translational modifications to histone and the proteins that read, write, and erase such modifications. The development of inhibitors for reader domains has focused on single domains. One of the major difficulties of designing inhibitors for reader domains, is that with the notable exception of bromodomains, they tend not to possess a well enclosed binding site amenable to small molecule inhibition. As many of the proteins in epigenetic regulation have multiple domains there are opportunities for designing inhibitors that bind at a domain-domain interface which provide a more suitable interaction pocket. Examination of X-ray structures of multiple domains involved in recognizing and modifying post-translational histone marks using the SiteMap algorithm identified potential binding sites at domain-domain interfaces. For the tandem plant homeodomain-bromodomain of SP100C, a potential inter-domain site identified computationally was validated experimentally by the discovery of ligands by X-ray crystallographic fragment screening.

## Introduction

Genetic information is contained in chromosomes which are made of chromatin, principally a combination of DNA and histone proteins. The repeating unit of chromatin is the nucleosome, which is composed of DNA wrapped around an octamer of histone proteins. In eukaryotes each nucleosome consists of two copies each of histone 2A (H2A), histone 2B (H2B), histone 3 (H3), and histone 4 (H4). N-Terminal histone tails protrude from the nucleosome and post-translational modification (PTM) of these tails control access to the nucleosomal DNA. PTMs of histone tails therefore play an important role in regulating gene expression. This process of epigenetic regulation is dynamic, and various writer-and eraser-enzyme families are involved in adding and removing post-translational modifications to histone tails. Although many different modifications of histones are known, acetylation and methylation of lysine found on the H3 and H4 tails are amongst the most extensively studied. The combination of PTM on histone proteins has been shown to be important in transcriptional regulation^[1]^ and has been termed the histone code. One aspect of the histone code hypothesis states that “distinct modifications of the histone tails induce different interaction affinities for chromatin-associated proteins.”

A second aspect of the hypothesis states that “modifications on the same or different histone tails may be interdependent and generate various combinations on any one nucleosome.” The predicted interdependence of histone tail modifications has been validated by the discovery of proteins that recognize multiple marks through multiple domains,^[2,3]^ and enzymes that require the presence of one mark in order to efficiently modify another.^[4]^ The ability for a protein to read multiple marks, or for an enzyme to create (write) or remove (erase) a mark based on the presence of another neighboring mark, probably requires multiple domains. Indeed, there are many examples of proteins that contain multiple domains known to bind and modify histone tails containing multiple PTMs (Figure 1).

**Figure 1.**
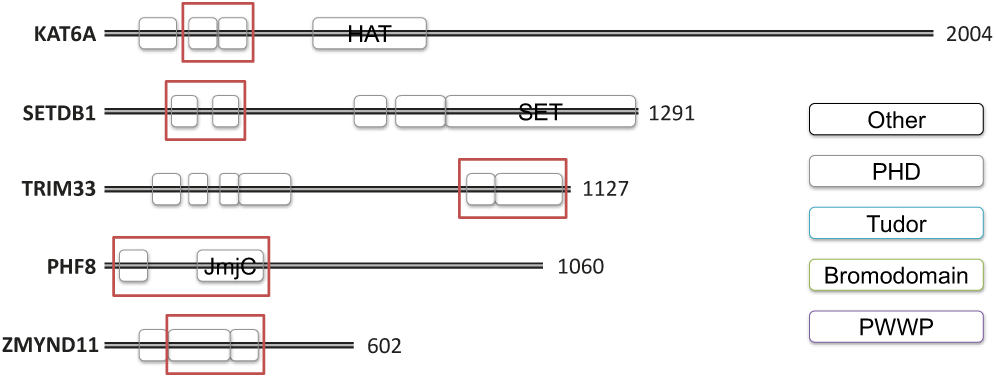
Domain maps exemplifying proteins that bind or modify histone tails containing multiple PTMs. Tandem domains studied in this work are in red boxes. HAT: histone acetyl transferase domain; KAT6A: lysine acetyl transferase 6; SETDB1: SET domain bifurcated 1; TRIM33: tripartite motif protein 33; PHF8: PHD Finger Protein 8; ZMYND11: Zinc finger MYND domain-containing protein 11; SET (Su(var)3-9, enhancer-of-zeste and trithorax): a lysine methyl transferase domain; JmjC (Jumonji C-terminal domain): an oxygenase lysine demethylase domain; PHD (plant homeo-domain), Tudor and PWWP are methyl lysine binding domains; bromodomains are acetyl lysine binding domains. Domain identity and boundaries are adapted from EBI Interpro.^[5]^

There have been many recent reports of inhibitors for reader,^[6– 15]^ writer,^[16–19]^ and, in some cases, eraser domains.^[20–24]^ Bromodomains (BRD) are acetyl-lysine binding domains, for which numerous inhibitors have been developed.^[25,26]^ Malignant Brain Tumour (MBT) domains are methyl-lysine binding domains and were the first methyl-lysine binding domains for which inhibitors were developed.^[13]^ Subsequently a peptidomimetic inhibitor has been developed for the methyl-lysine binding chromodomain of chromobox homologue 7 (CBX7),^[12]^ as well as a small molecule inhibitor of the third PHD of KDM5A^[11]^ although both of these examples have low potency in the micromolar range. Inhibitors have also been developed for both methyltransferases/demethylases^[16]^ and histone acetyltransferases/deacetylases (HATs/ HDACs).^[17,27]^

As well as the development of inhibitors for pharmaceutical use, there have been a number of inhibitors designed to be used as chemical probes.^[28,29]^ These are tool compounds with proven in cell target engagement and suitable selectivity and potency that can be used for target investigation and validation experiments (Figure 2).

**Figure 2.**
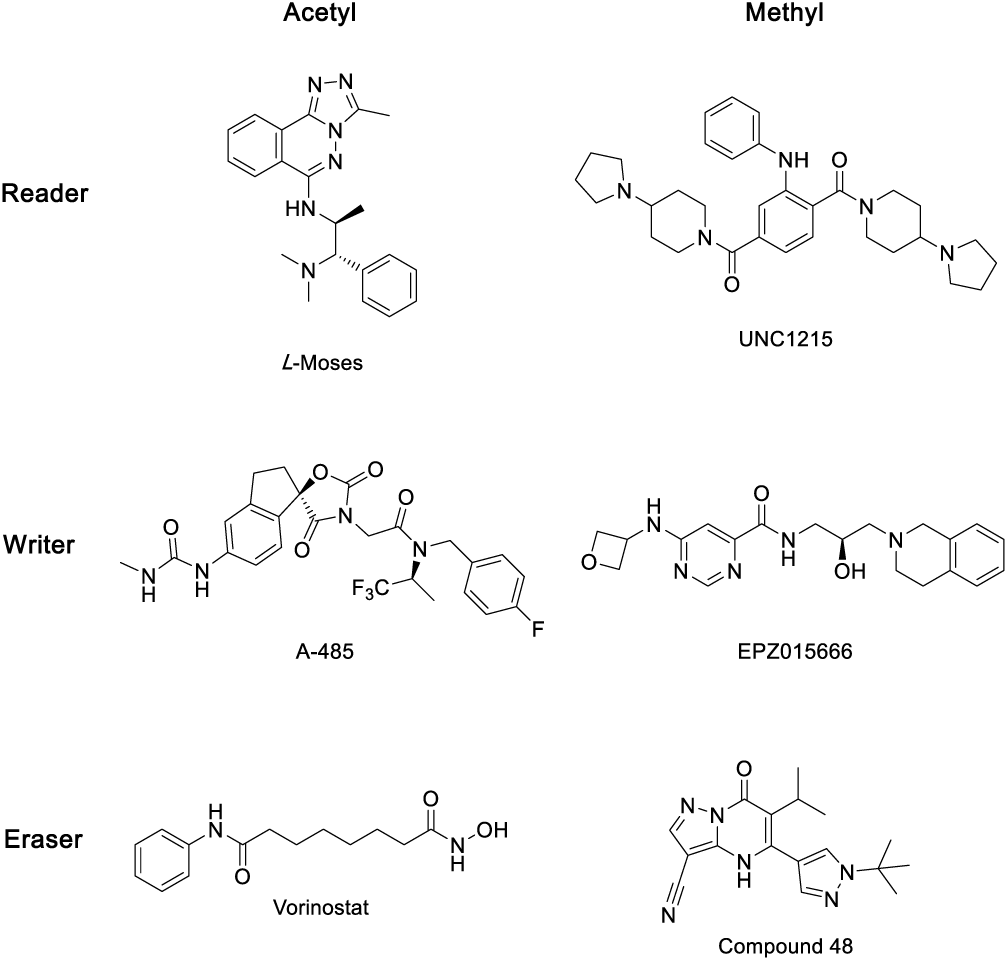
Exemplary inhibitors of readers, writers, and erasers. *L*-Moses^[15]^ and UNC1215^[13]^ are inhibitors of the BRD of PCAF and methyl-lysine binding MBT domain of L3MBTL3 respectively. A-485^[19]^ is a CBP/p300 histone acetyl transferase (HAT) inhibitor, and EPZ015666^[16]^ inhibits the methyl transferases PRMT5. Vorinostat (SAHA) is a clinically used HDAC inhibitor and compound 48^[24]^ is an inhibitor of the 2-oxoglutarate dependent KDM5 demethylases.

Despite these successes, there are still many families of protein domains involved in reading, writing, or erasing histone tail modifications with very poor inhibitor coverage. Of these many are predicted to be targets with intrinsically low ligandability.^[30]^ A study by Santiago *et al.*^[31]^ suggested that of methyl-lysine readers, MBT domains, Tudor domains, and PHDs are inherently less ‘ligandable’ than other epigenetic reader domains. During our investigations into inhibitors of PHDs, we observed that tandem PHDs appear more druggable with deeper and more hydrophobic pockets than isolated single PHDs and we postulated that binding sites may exist between adjacent domains in other epigenetic families that would be more amenable to ligand discovery than the individual domains considered in isolation.

There are increasing reports of X-ray crystal structures of multiple domains involved in recognizing and modifying post-translational marks on histone tails. These include examples of multiple reader domains from the same protein,^[32,33]^ and also reader-writer^[34]^ and reader-eraser combinations.^[4]^ Structures containing multiple epigenetic domains provide us with an opportunity to study sites formed at the domain-domain interfaces, and investigate whether these sites are suitable for ligand binding.

## Results and Discussion

We used SiteMap^[35]^ to analyze structures containing multiple histone binding/modifying domains in order to identify potential ligand binding sites at domain-domain interfaces. It is hoped that these sites offer a potential solution for inhibitor design of histone binding/modifying proteins, where the individual domains possess sites with low ligandability.

SiteMap works by creating a grid of points separated by 1 Å. Each point is then evaluated to determine whether it overlaps with any protein atoms, points that overlap with protein atoms are removed. Further points are removed if they are insufficiently enclosed by the protein or if they have a too small Van der Waals interaction energy with the protein. Points are then grouped into sites, and the enclosure, size, and hydrophilicity of the site calculated. These values are combined into a single scoring function called the SiteScore.^[36]^

### Prevalence and Ligandability of multi-domains

The structures used in this study were identified using ChromoHub.^[37,38]^ Any structure of a histone reader, writer, or eraser which also contained another histone reader, writer, or eraser domain was included. A total of 103 structures were identified, representing thirty-three unique gene products (Figure 3). The most common domain combination was multiple-Tudor domains of which seven examples were identified.

**Figure 3.**
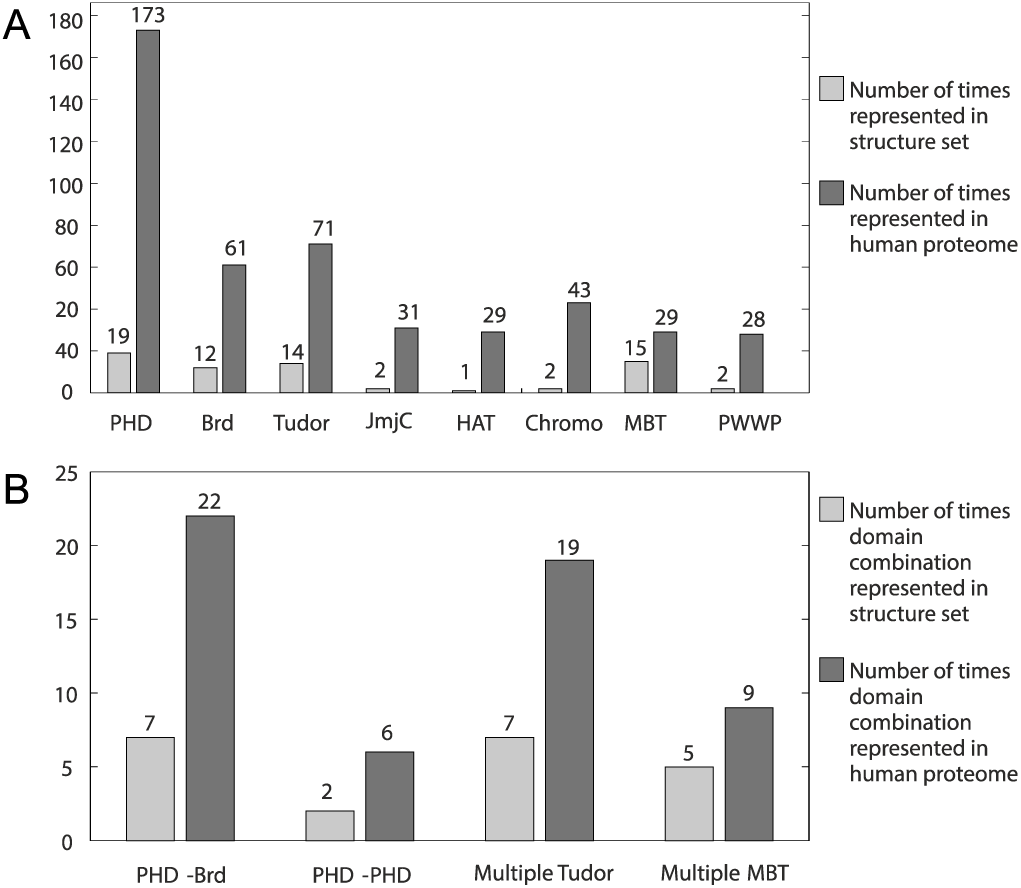
Occurrences of epigenetic domains in tandem with other domain structures reported in the PDB A. PHDs, BRDs, Tudor domains, and MBT domains are the most common domains in the tandem structure set. B. Structures containing multiple Tudor domains appear seven times, with five structures containing multiple MBT domains. The most common hetero-domain combination in our structure set is PHD-BRD, of which seven examples were identified. A PHD-BRD has an N-terminal PHD and a C-terminal BRD; this is distinct from a BRD-PHD, in which the domain order is reversed.

The domain combinations in Figure 3B are all adjacent to each other in the protein sequence, or have crystallographic evidence that they act together to engage histones. Domain combinations can have the histone binding potential greatly affected by the linker between the two domains. It is interesting to compare the JmjC-PHD domain combination found in KDM7B (PHF8) with the JmjC-PHD domain combination found in KDM7A (KIA1718). The two JmjC domains, and the two PHDs are very closely homologous (63% and 77% sequence identity respectively); however, differences in the linker (which is much longer in KDM7A) effect the relative conformation of the two domains and hence the effects of the PHD domains on the specificity of the demethylase activity of the JmjC domain.^[4]^

SiteMap was used to calculate a SiteScore for every multidomain combination identified. In general it was possible to identify potential binding sites in most structures, and in some cases these were found at domain-domain interfaces (Figure 4). Specific examples of ligandable tandem domains are discussed in detail below.

**Figure 4.**
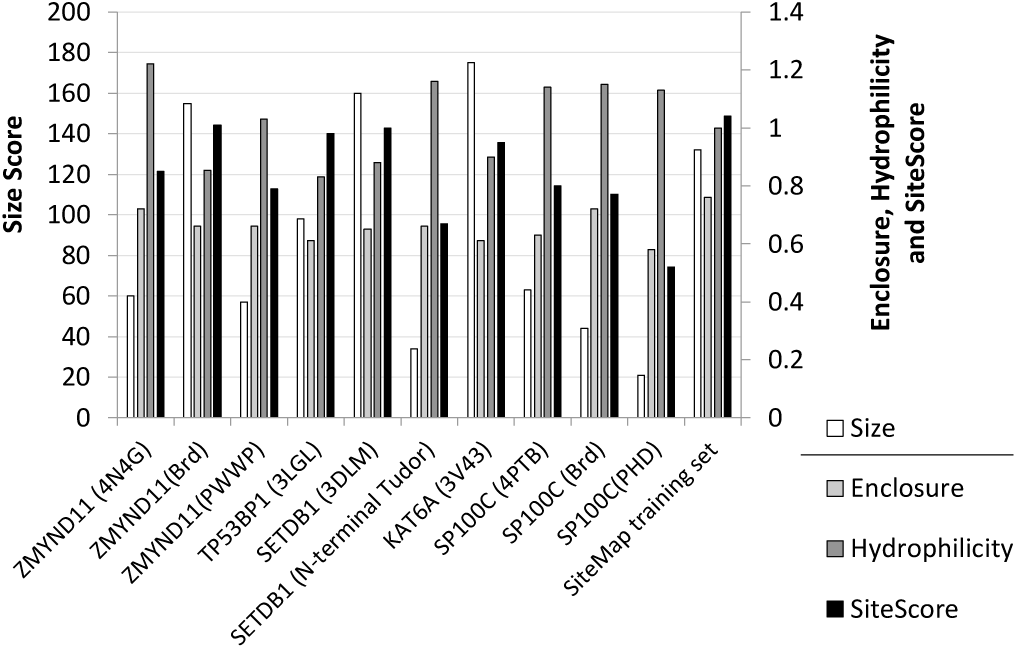
The size, enclosure, hydrophilicity, and SiteScore of selected tandem domains. The overall SiteScore is a combination of the sites size, enclosure and hydrophilicity. The mean values for the 326 binding sites with known submicromolar ligands that were used as a training set during the development of SiteMap is shown to benchmark the analysis of the tandem domains.

### Bromodomain-PWWPs

The structure of the BRD-PWWP domain of tumour suppressor ZMYND11 has been solved with a bound H3.3 peptide (Figure 5).^[39]^ Wen et al. identified an interaction between a serine residue (S31) found in H3.3 and the BRD-PWWP domain-domain interface. This recognition allows ZMYND11 to differentiate between the H3.3 variant and the more common H3 histone, which has an alanine at position 31. Isothermal titration calorimetry (ITC) reveals a 7-fold difference in binding to ZMYND11 between H3 tri-methylated on lysine-36 (H3K36me3 K_*D*_ 431 µM) and a K36-trimethylated H3.3 peptide, residues 19-42 (H3.3_19-42_K36me3 K_*D*_ 56 µM).^[39]^ Analysis of the structure of the BRD-PWWP of ZMYND11 (PDB ID 4N4G) with SiteMap identified the BRD as the most ligandable site (SiteScore = 1.01). However Wen *et al.* described this BRD as “unlikely to be a histone acetyl-lysine-binding module” due to the presence of a tyrosine residue at the position where asparagine is commonly found in acetyl-lysine binding BRDs. ^[39]^ Therefore the BRD offers limited potential for the development of a small-molecule inhibitor of the histone binding function of ZMYND11.

**Figure 5.**
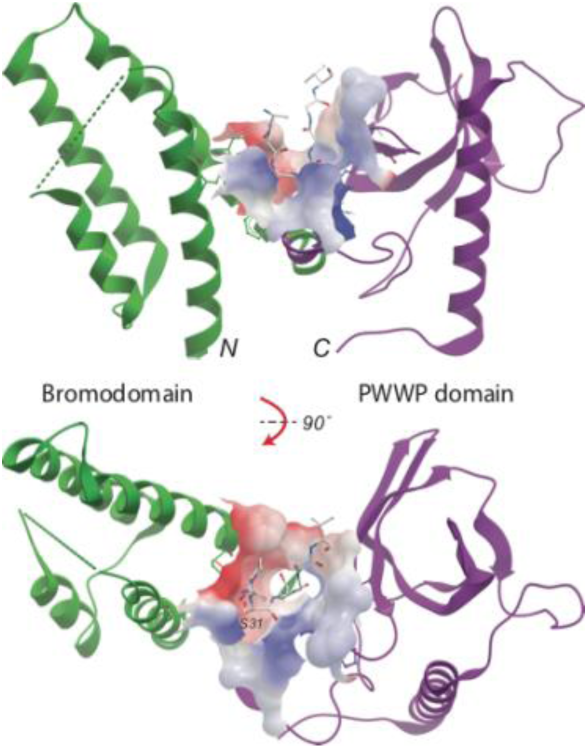
View of a crystal structure of the BRD (left, green) and the PWWP domain (right, magenta) of ZMYND11 (PDB ID 4N4G). Residues 29-39 of H3.3 are shown in stick form at the histone binding site identified between the two domains (surface).

A binding site at the BRD-PWWP interface where H3.3 S31 binds is identified as having a SiteScore of 0.85. This is at the lower end of what would be considered ligandable.^[36]^ Further analysis of this binding site shows it is a well enclosed site, with a SiteMap enclosure score of 0.72, this compares favorably to the mean value 0.76 for the sub-micromolar sites in SiteMap’s training set. The site is relatively small; with a SiteMap size score of 60, whereas the mean size score of the sub-micromolar sites in the training set is 132. However, the relatively smaller size at the BRD-PWWP interface should not necessarily be inhibitive for ligand design. Chemical probe discovery efforts directed at the H3.3 S31 binding site is more likely to deliver a potent inhibitor than a single-domain inhibitor of the BRD or PWWP domains.

### Tudor domains

Our structure set contains seven Tudor domain containing proteins. All seven of these examples contain a multiple Tudor domain, with one example (UHRF1) containing a tandem Tudor and a PHD. In most of these cases, only one of the multiple Tudor domains contains the arrangement of aromatic side chains commonly known as an aromatic cage that is typically found in Tudor domains and plays a crucial role in binding to methylated lysines.^[40]^ Analysis of a series of Tudor domains (both isolated and tandems) showed that the aromatic cage typically provides a well enclosed hydrophobic region that could be used to anchor an inhibitor. However, these enclosed regions are typically small, with size scores typically less than 40, hence these sites tend to have low SiteScores. In the case of the tandem Tudor domains, the second domain provides some expansion space around the aromatic cage. This creates a site where an anchoring head group could bind to the aromatic cage, with the rest of the molecule interacting with the expanded surface created by the second Tudor domain and methyl-lysine mimetics could exploit these aromatic cages, in a similar way to how acetyl-lysine mimetics have been used to produce BRD inhibitors.^[10]^

The tandem Tudor domain of DNA repair factor TP53BP1 (tumour protein p53 binding protein 1) is an illustrative example of the case described above. Our analysis by SiteMap reveals a site with a SiteScore of 0.98 containing the aromatic cage of *N*-terminal Tudor and extending to an area at the interface of the two Tudor domains, (Figure 6A). This site is large and open, with size and enclosure scores of 98 and 0.61 respectively. Although this is a very open site, it is amenable to techniques designed for the discovery of inhibitors of protein-protein interactions and it would be expected that due to its incorporation of the aromatic Tudor cage, that ligands binding at this tandem site would interfere with TDP53BP1 substrate binding.^[41]^

**Figure 6.**
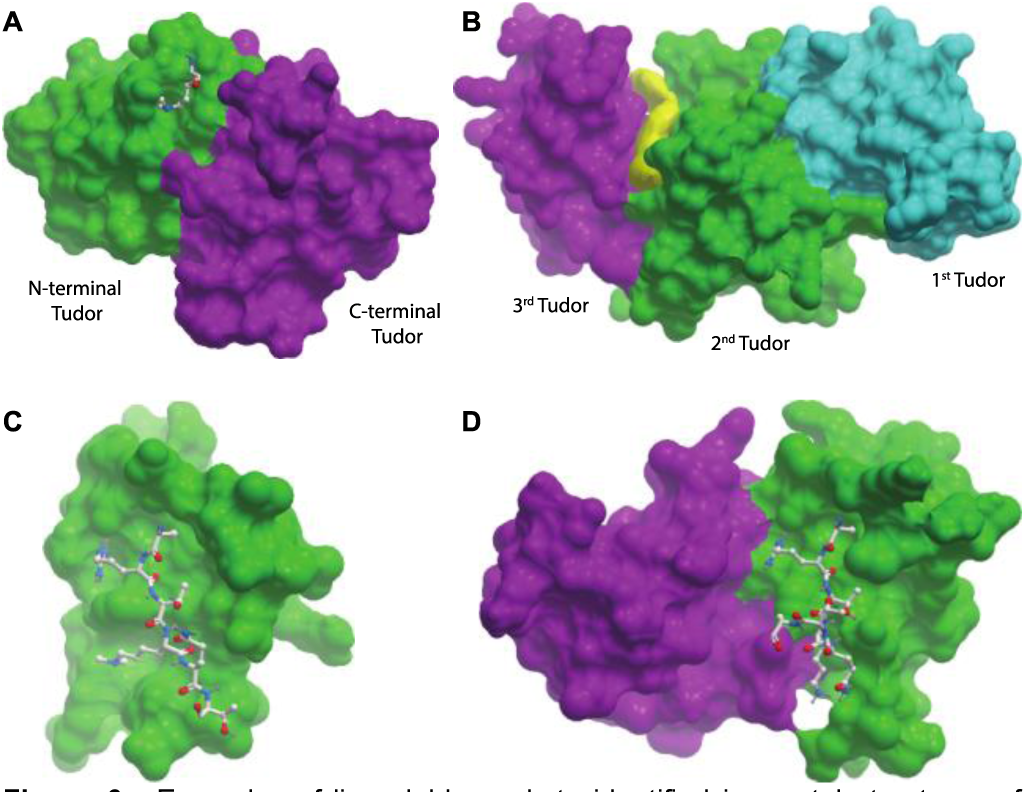
Examples of ligandable pockets identified in crystal structures of PHD and Tudor domain proteins. A. The H3K4me2 residue is present in the binding site of the *N*-terminal Tudor (green) of TP53BP1, near the interface with the *C*-terminal Tudor domain (magenta) (PDB ID 3LGL). B. Yellow coloring indicates the volume of the ligandable cleft between the second (green) and third (magenta) Tudor domains of SETDB1 is (PDB ID 3DLM). C. The single PHD shown is that of BPTF (PDB ID 2FSA). D. the tandem PHD shown is that of KAT6A (PDB ID 3V43). The *N*-terminal PHD of KAT6A is magenta and the *C*-terminal PHD is green.

The histone methyltransferase SETDB1 contains three contiguous Tudor domains. The available crystal structure (PDB ID 3DLM) does not show a well formed aromatic cage in any of the three Tudor domains. However, closer inspection reveals that the second and third Tudor domains contain suitable residues to form an aromatic cage, the formation of which could be induced by binding of a tri-methylated lysine. The first Tudor domain also contains two aromatic residues but this potential aromatic cage is blocked by a lysine side chain. The triple Tudor domain contains a potential ligand binding site between the second and third Tudor domain (Figure 6B). As discussed above, these are the two most likely to be involved in binding to a *N*-methyl-lysine residue, and therefore a ligand that binds between the second and third Tudors would impair peptide binding. This site has a SiteScore of 1.00 and this good score is primarily down to the site’s large size (size score of 160) and low hydrophilicity (0.88).

### PHD domains

It has been predicted that PHDs in general have low ligandability, however there are some exceptions. One example of a ligandable site at a domain-domain interface involving a PHD has already been identified.^[31]^ The demethylase KDM7B (PHF8) contains a cavity at the interface of its PHD and JmjC domains. This site is involved in histone binding, and is therefore of interest for the design of a substrate competitive KDM7B inhibitor.

Our structure set contains three examples (DPF3, MLL3, and KAT6A) of PHD-PHDs. The structures of DPF3 and KAT6A show that the tandem PHD module interacts with residues 1-14 of H3, with H3 binding at the domain-domain interface. Tandem PHDs have been shown to be acetyl-lysine binders; this acetyl-lysine recognition takes place at a site formed by the domain-domain interface.^[42]^ SiteMap analysis of the H3 binding site of acetyl transferase KAT6A (MYST3, PDB ID 3V43) identifies the H3 binding site as having a site score of 0.95. Although this site is large (size score 175) it is as well quite open, with an enclosure score of 0.61. Although it would be considered difficult to design a conventional ligand for this type of pocket, this pocket would be a suitable candidate for techniques designed to target protein-protein interactions.^[41]^ KAT6A is known to form fusion proteins containing the tandem PHD region with the histone acetyl transferases CREBBP and p300, forming an aberrant acetylation complex that plays a role in acute myeloid leukaemia.^[43]^ Selective inhibition of the interaction of the tandem PHDs of KAT6A with chromatin would allow study of the role these domains play in the protein fusion and in the disease progression. SiteMap analysis of the PHD-PHD of BAF chromatin remodelling complex member DPF3 shows that it also contains a site at the domain-domain interface that is more promising than a typical site of a single PHD. SiteMap analysis of the 20 NMR models from the PDB deposition 2KWJ identifies a site with a mean SiteScore of 0.81. Similarly to KAT6A, the site identified in DPF3 is large and open, with a mean size score of 72 and a mean enclosure of 0.59. This suggests that a PHD-PHD is more ligandable target than a single PHD (Figure 6C,D).

There are 20 structures containing a PHD and another domain in our structure set. The most common partner domain being a *C*-terminal BRD (8 examples). PHD-BRDs have been shown to act together in substrate recognition.^[2]^ The PHD-BRD structures investigated in this study fell into two categories, those where the PHD and BRD form a compact globular structure, and those where the PHD and BRD are separated by a rigid linker (Figure 7).

**Figure 7.**
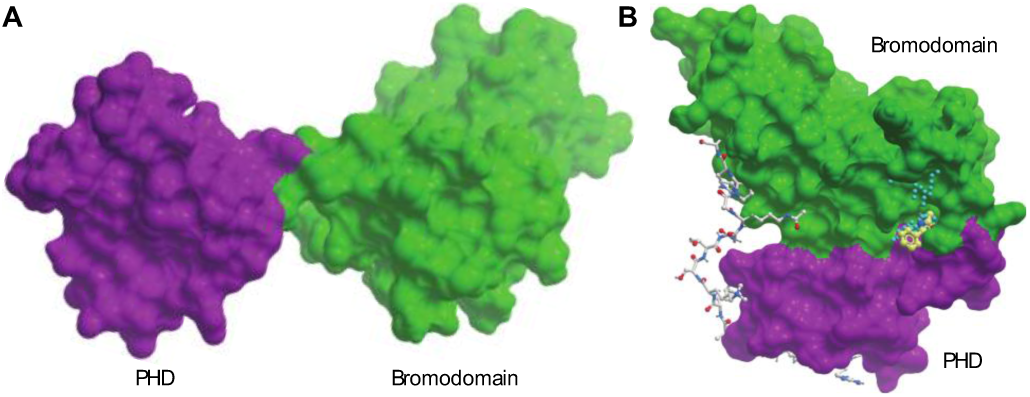
Examples of two types of PHD-bromodomain. A. The PHD domain (magenta) and BRD domain (green) of BPTF (PDB ID 2FSA) are separated by a rigid linker. B. The PHD domain (magenta) and BRD domain (green) of SP100C (PDB ID 4PTB) form a compact, globular structure. SP100C is shown with the site points as identified by SiteMap, and an overlaid ligand identified by a crystal soaking experiment. The novel potential ligand binding site at the domain-domain interface of the PHD-BRD of SP100C is indicated by cyan dots, the PHD is shown in magenta, and the BRD is shown in green. A peptide taken from a structure of the related PHD-BRD of TRIM33 (PDB ID 3U5P) is superimposed to highlight the probable peptide binding surface.

Amongst the group of structures with a globular arrangement of the PHD and BRD is SP100C. Analysis of this structure by SiteMap reveals the presence of a novel potential ligand binding site at the interface of the PHD and BRD domains (Figure 7B). This site has a SiteScore of 0.81, which would place it in the range of challenging but ligandable binding sites. Further analysis of the site reveals that it is large and well enclosed, but suffers a penalty to its SiteScore due to its hydrophilic nature.

### SP100C Fragment Screening

In order to further investigate this newly identified potential binding site of SP100C, we pursued structural and ligand binding studies on the tandem BRD-PHD construct.^[44]^ X-ray crystallography experiment was performed at the XChem facility of Diamond Light Source where SP100C crystals were soaked with 412 individual fragment compounds at 30 mM.^[45]^ In addition to the typical weak protein surface binders, this soaking experiment revealed only two high confidence ligands, both of which bind in the novel pocket at the interface of the PHD and BRD domains (Figure 8). Although the binding affinity of these compounds was not determined, they were clearly present in the crystal structure and surprisingly no fragments were found bound at the single-domain BRD or PHD pockets. Based on comparisons to the related PHD-BRD of TRIM33,^[2]^ this pocket is not expected to be involved in histone peptide recognition. However, it is possible that a ligand binding at this position could allosterically modify the PHD-BRD interface.

**Figure 8.**
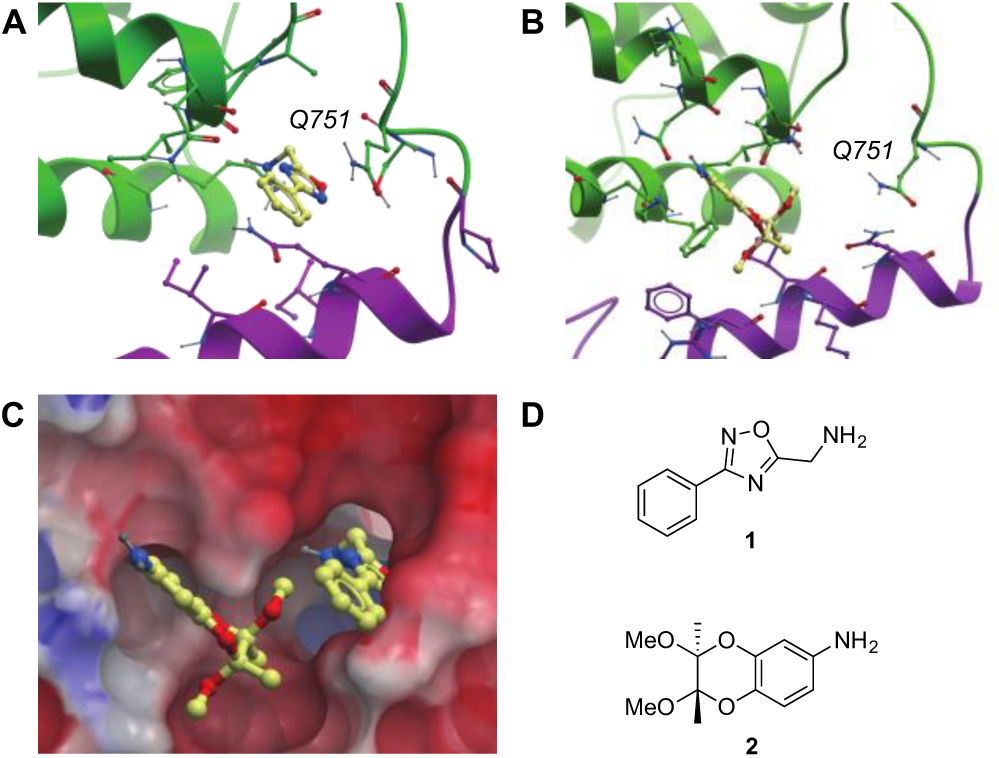
Fragment soaking in SP100C identifies compounds binding at the PHD/BRD domain interface. A. Binding mode of oxadiazole **1** (yellow) to SP100C. The backbone ribbon and side chains of residues from the PHD domain involved in the compound binding are colored in magenta, the backbone ribbon and residues from the BRD domain involved in the compound binding are colored in green. B. Aniline **2** does not bind as deeply in the pocket as oxadiazole 1 and therefore does not cause a conformational change in residue Q751. C. The two fragment hits identified for the inter-domain binding site of SP100C are shown overlaid with an electrostatic map of the surface of the *apo* structure of SP100C. Note that the SP100C inter domain binding site presents a more hydrophobic surface than the rest of the surface area. D. Chemical structures of compounds **1** and **2**.

Oxadiazole **1** binds deeper into the pocket than aniline **2**. It exploits hydrophobic interactions with surrounding residues. Oxadiazole **1** is positioned to form a hydrogen bond with the backbone carbonyl of residue I871 via its primary amine (Figure 8A). It also induces a conformational change in residue Q751 not observed for aniline **2**. The primary amide of Q751 is rotated through 120° relative to the uncomplexed structure. Aniline **2**^[46]^ does not bind as deeply in the pocket as oxadiazole **1**, and therefore does not cause any movement of residue Q751 relative to the *apo* structure (Figure 8B). It also exploits hydrophobic interactions with surrounding residues. A comparison of the binding mode of the two identified fragments shows that oxadiazole **1** binds deeply into the pocket predicted by SiteMap, with aniline **2** occupying a hydrophobic site at the mouth of the pocket. This arrangement of these hits suggests that a fragment linking strategy could lead to optimized ligands (Figure 8).

## Conclusions

Developing inhibitors of epigenetic proteins is of growing interest in the quest for new treatments for diseases such as cancer and inflammation. Inhibitors have been developed for some domain families involved in histone binding and modification. So far three inhibitors for HDACs are in clinical use.^[47]^ Despite these successes, there are many domain families, especially the reader domains, for which few inhibitors have been discovered. As many epigenetic proteins have tandem reader domains, targeting cavities formed at domain-domain interfaces offers an alternative to develop inhibitors rather than targeting individual reader domains only. We have shown that tandem reader domains often present a more ligandable pocket either due to a larger inter domain area (PHD-PHD) or to a unique pocket located at the domain interface (SP100C).

In the case of the PHD-BRD tandem domain of SP100C, we have identified a novel binding site using SiteMap, and identified two fragments that bind at this position by performing a X-ray fragment screening campaign at the Diamond synchrotron light source. This crystallographic evidence validates the predictions made by SiteMap in the case of SP100C. It also provides an example of a method by which initial hits could be identified for the other identified inter-domain binding sites.

In the case of PHD-PHDs, and the BRD-PWWP of ZMYND11 the identified inter-domain binding sites appear to play an important role in histone tail recognition. The potential ligand binding site identified on ZMYND11 is directly involved in the discrimination between H3.3 and the more common H3.1. Therefore, as well as being a suitable binding site for a small molecule ligand, it also plays an important role biologically. Similarly, the binding site identified at the PHD-PHD interface in KAT6A is both a promising binding site for a small molecule ligand and of biological relevance as an acetyl-lysine binder. These examples show that inter-domain binding sites have substantial potential for the development of inhibitors of protein-histone interactions which could yield useful chemical probes for the study of protein complexes involved in epigenetic diseases.

## Experimental Section

### Computational Analysis of Binding Sites

The structures used in this study were identified using ChromoHub.^[37,38]^ Any structure of a histone reader, writer, or eraser which also contained another histone reader, writer, or eraser domain was included. A total of 103 structures were identified, representing thirty-three unique gene products. Schrödinger SiteMap (version 2.8, Schrödinger, LLC, New York, NY, 2013) was run from the command line using a Linux server. Structures were imported into Schrödinger Maestro in pdb file format. Additional chains, domains, and water molecules were removed. The structures were process using the Schrödinger Protein Preparation Wizard, and bound ligands removed. SiteMap was run using default parameters unless otherwise specified.

### SP100C Protein Crystallography and Fragment Soaking

SP100C crystals were grown in SWISSCI 3 Lens crystallization sitting-drop plates at 4 °C by mixing 50–100 nl of 10 mg/ml protein solution in a 1:1 ratio with 50–100 nl reservoir solution consisting of 0.1 M MES pH 7.0, 20–30% (w/v) PEG 20000 and placing the drops over 20 μL reservoir solution. Crystals appeared in 6-7 days. Crystal soaking was performed by liquid droplet transfer using a TTPLabtech Mosquito® HTS. Ethylene glycol was added for cryoprotection using the Mosquito® to a final concentration of 25% (v/v) (calculated from the initial drop volume). The SP100C crystals diffracted to 1.6–2.0 Å resolution in space group C2, with typical unit-cell parameters a=127.4Å, b=45.4Å, c=84.0Å, β=102.0° and with one SP100C molecule in the asymmetric unit.

X-ray diffraction data were collected on beamline I04-1 at Diamond Light Source and were processed using the Diamond autoprocessing pipeline, which utilizes xia2,^[48]^ DIALS,^[49]^ XDS,^[50]^ POINTLESS^[51]^ and CCP4.^[52]^ Electron-density maps were generated using XChemExplorer^[45]^ via DIMPLE.^[53]^ Ligand restraints were generated with AceDRG^[54]^and iterative refinement and manual model correction was performed using REFMAC^[55]^ and Coot,^[56]^ respectively.

## Accession Codes

Atomic coordinates and structure factors for the crystal structures of SP100C with compounds **1** and **2** can be accessed using PDB codes XXXX and XXXX, respectively. Authors will release the atomic coordinates and experimental data upon article publication.

## Acknowledgements

The SGC is a registered charity (number 1097737) that receives funds from AbbVie, Bayer Pharma AG, Boehringer Ingelheim, Canada Foundation for Innovation, Eshelman Institute for Innovation, Genome Canada, Innovative Medicines Initiative (EU/EFPIA) [ULTRA-DD grant no. 115766], Janssen, Merck KGaA Darmstadt Germany, MSD, Novartis Pharma AG, Ontario Ministry of Economic Development and Innovation, Pfizer, Sao Paulo Research Foundation-FAPESP, Takeda, and Wellcome [106169/ZZ14/Z]. We are grateful to Chandra Ramarao of Avra Laboratories and 5+1C for the generous gift of compound **2** for our fragment collection.

## Entry for the Table of Contents

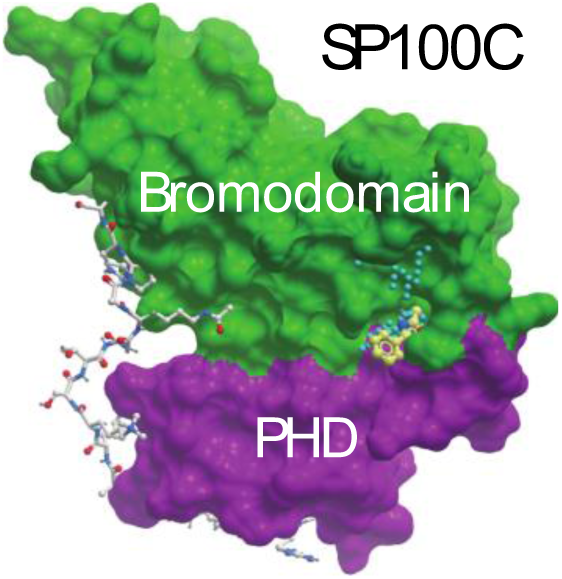

Many epigenetic proteins have adjacent domains that bind multiple histone PTMs. Examination of structures of these tandem domains identified binding sites at the interdomain interface which often have better ligand binding properties than their respective single domains. For the tandem PHD-bromodomain of SP100C, an interdomain site was validated experimentally by the discovery of fragment ligands by X-ray crystallographic screening.

